# There is no Biophysical Distinction between Temporal Interference and Direct kHz Stimulation of Peripheral Nerves

**DOI:** 10.1101/2024.09.06.611584

**Authors:** Aleksandar Opančar, Petra Ondráčková, David Samuel Rose, Jan Trajlinek, Vedran Đerek, Eric Daniel Głowacki

## Abstract

Temporal interference stimulation (TIS) has attracted increasing attention as a promising noninvasive electrical stimulation method. Despite positive results and optimistic expectations, the TIS field has been beset by misunderstandings concerning its mechanism of action and efficacy in safely targeting deep neural structures. Various studies posit that TIS exploits the low-frequency “beat” frequency produced by interference of multiple kHz carriers to essentially deliver low-frequency stimulation at the intersection of the carriers, thereby circumventing limitations associated with tissue impedance and depth penetration. Due to the documented electrophysiological effects of kHz-range stimuli, such a picture is an oversimplification, and at present a critical open question for TIS is how much stimulation thresholds practically differ for amplitude-modulated versus unmodulated kHz stimuli. This paper presents experimental evidence supporting the conclusion that TIS is driven by the kHz carrier itself, and that amplitude modulation frequency has little relatively effect on thresholds. We test TIS and different modulated/unmodulated kHz waveforms on peripheral nerve targets in an invertebrate model (*Locusta migratoria*), and on sensory and motor stimulation in human subjects. We find that modulated (TIS) and unmodulated sine carrier frequencies in the range 0.5 – 12.5 kHz in humans, and further up to 100 kHz in locusts, demonstrate overlapping strength-frequency (s-f) dependence across all stimulation types. Since all stimulation waveforms are governed by the same s-f curve, this implicates a common underlying biophysical mechanism. This equivalence challenges the notion that the TIS modulated “envelope” frequency can uniquely facilitate neural engagement via other mechanisms. We test amplitude-modulation frequency (AMF) dependence, confirming the resonance effect predicted by kHz rectification theory, and we evaluate the regions of tonic (unmodulated) and phasic (amplitude-modulated) stimulation regions inherent when using TIS. Our results help to resolve the mechanistic debate, at least for suprathreshold TIS. We explain how the findings can be extrapolated to subthreshold stimulation, and further suggest possible advantages of using 2-electrode amplitude-modulated kHz waveforms over the multielectrode TIS configurations.

## 1. Introduction

### Noninvasive electrical stimulation and the use of interferential currents

With electrical neurostimulation of the nervous system there is always the tradeoff between invasiveness and specificity. Applying small electrodes to the exact area of interest is possible with high precision, however at the cost of a complex, invasive, and to some degree damaging surgical procedure^1–3^. Completely noninvasive electrical stimulation from outside the body is much easier to perform, however it is plagued by low spatial specificity, and limited depth of effective penetration^4^. This means only superficial targets can be effectively stimulated, for instance nerves located near the skin surface, or brain structures directly below the skull. Obtaining suprathreshold neurostimulation, where neurons become sufficiently depolarized and directly fire action potentials, requires electric fields (> 1V/m) which are difficult to obtain transcutaneously without causing discomfort arising from stimulation of sensory receptors and fibers in superficial tissues.^5,6^ This greatly limits the diagnostic and therapeutic potential of noninvasive electrical stimulation, and implantable electrodes remain necessary for many applications where direct triggering of action potentials is desired (*i.e.* suprathreshold neurostimulation). The fundamental limitation behind noninvasive central or peripheral electrical stimulation is that electric field decreases inversely with distance from the electrodes for any given conductive medium. The relatively high impedance of skin imposes practical limits to penetration of electrical field^6^. The challenge of how to effectively deliver electrical stimuli through the skin is a fundamental problem for noninvasive bioelectronic medicine. A suggested approach to overcome this basic issue is using high-frequency electric fields, since skin impedance decreases with frequency^7^. The higher the carrier frequency used, the lower is the effective impedance. One method to exploit this, which this paper interrogates, is the concept of mixing multiple high-frequency electrical signals to produce patterns of interference and amplitude-modulated (AM) “beats” in the tissue. This amplitude modulation frequency (AMF) is assumed to be the effective stimulation frequency recruiting neural firing. This interference method could solve the issue of depth penetration and potentially spatial specificity also. This idea was originally suggested by Nemec in the 1950s under the name “interferential current stimulation” (ICS). Since then, ICS has been widely adopted in the field of peripheral nerve stimulation, for instance in functional electrical stimulation, physiotherapy, and sports medicine^8–13^. The kHz interference method has gained increasing attention since its demonstration for brain stimulation in mice in the 2017 paper by Grossman *et al.*.^14^ While in the older literature the terms ICS or “interferential therapy, “ICT” are more common, in recent studies, this type of stimulation using multiple kHz frequency carriers is termed temporal interference stimulation (TIS)^15–20^. TIS has shown efficacy in humans for stimulation of peripheral nerves^21^, and promising examples of its use in brain stimulation have been reported^20,22^. In the literature, studies employing the term ICS typically involve suprathreshold stimulation of peripheral nerve targets, while TIS is the term more often associated with brain stimulation experiments, though in recent years “TIS” often replaces “ICS” for peripheral nerve studies as well. The terms ICS and TIS are both used in the literature since 2017. For clarity, we will use the term TIS in the remainder of this manuscript.

The principle behind TIS (Figure 1A) is that relatively high-frequency electrical stimulation signals, known as carriers, are applied, in a range over 1000 Hz. In TIS, it is assumed that these > 1 kHz stimuli do not efficiently recruit electrophysiological activity on their own. Many papers describing TIS assume *a priori* that the carriers are of too high frequency and too low amplitude to efficiently elicit action potential firing by themselves^14,16,19,20,23^. To accomplish TIS, multiple carriers of slightly different frequencies (*i.e. f*1 and *f*2) are applied simultaneously. This gives an offset between the two frequencies, Δ*f* (Figure 1A). The two carriers interact inside of the tissue, coming in and out of phase with each other, a phenomenon known as interference. Interference causes amplitude modulation (AM) of the resultant kHz electric fields inside the tissue. The amplitude modulation frequency (AMF) is equal to Δ*f*. The two frequencies *f*1 and *f*2 thus create an AM “envelope” frequency at Δ*f*. A number of published TIS studies establish that suprathreshold electrical neurostimulation synchronous to the AMF is indeed occurring^14,21,24,25^. This has led to two competing interpretations: A) the AMF produced by the interference effectively causes phasic stimulation at the target site because of intrinsic preference of neurons for lower frequencies, while the unmodulated high-frequency carrier has a negligible effect on surrounding tissues. In other words, the threshold for AM stimulation is significantly lower than the threshold of stimulation by the unmodulated carrier^14,26^; and neurons demodulate the AMF. B) TIS leads to phasic stimulation in regions of interference producing AM, while other regions are subject to equal or higher degrees of tonic stimulation by the carrier frequency, since biophysically the origin of the phasic and tonic stimulation is the same, and no selective demodulation of the AMF exists. Interpretation B was explained in a theory paper by Mirzakhalili et al.^17^. This paper motivated us to test these hypotheses in practice: we compare the stimulation thresholds of different modulated and unmodulated kHz waveforms (Figure 1B). Namely, we test a 4-electrode TIS configurations versus a 2-electrode AM-sine (these two conditions produce equivalent modulated E-fields), an on/off modulated sine (Sine burst), and finally an unmodulated sine. We measure these four conditions as a function of carrier frequency, the strength-frequency, s-f, relation (0.5 kHz – 12.5 kHz) in insect and human peripheral nerve models (Figure 1C). We find qualitatively similar stimulation using all waveforms, and essentially overlapping s-f relations of TIS and other kHz waveforms. Taken together, our results support that interpretation B is the correct.

**Figure 1.**
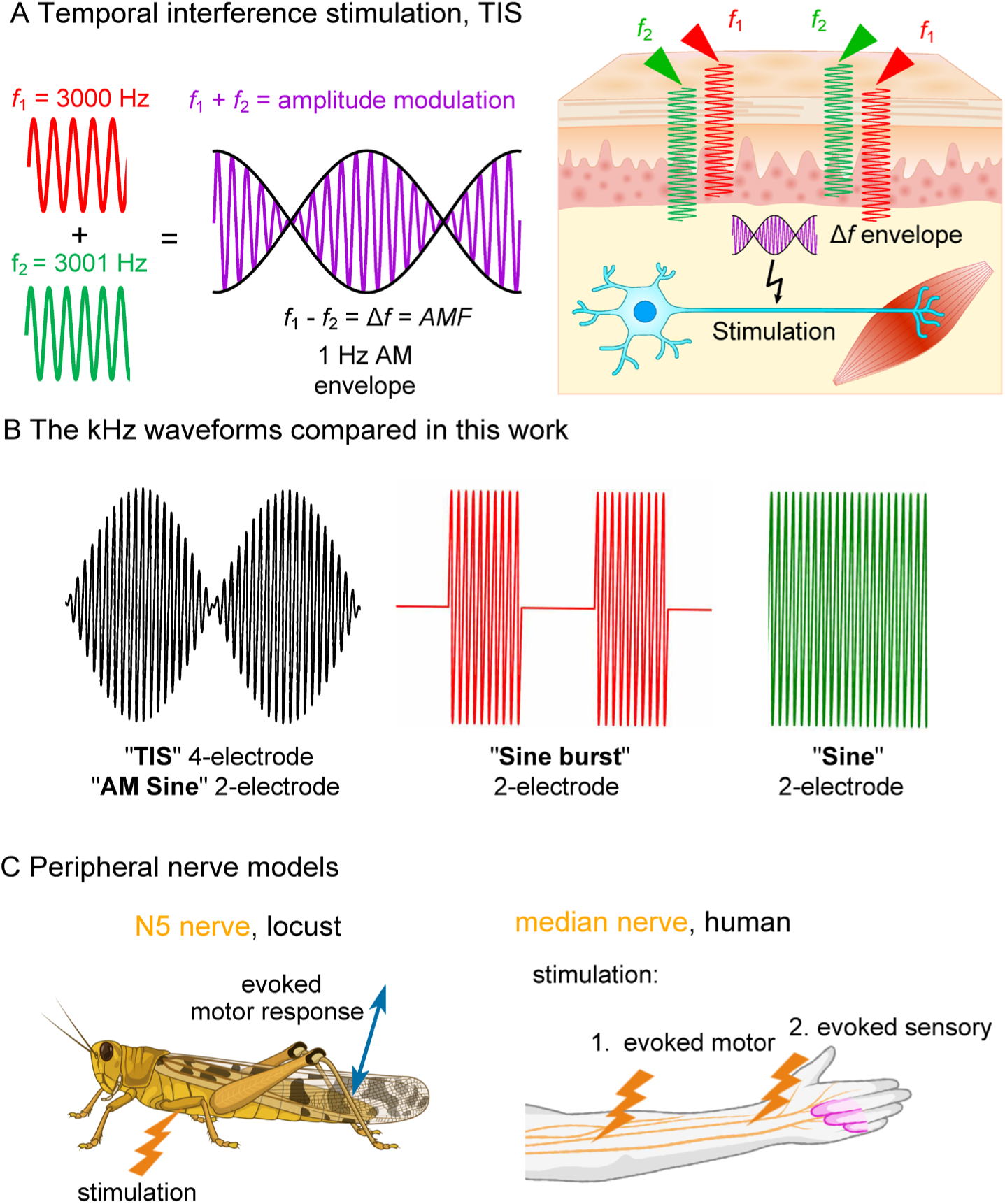
Testing the assumptions of TIS with respect to kHz electrical stimulation. A, The principle of TIS involves two kHz carriers at two different frequencies, *f*1 and *f*2, which interact in a given medium via interference, creating areas of amplitude modulation (AM). Maximum AM depth is achieved at midpoints between stimulation electrodes. The AM occurs at a frequency which will be equal to the frequency difference between the two carriers, Δ*f.* TIS can be used *in vivo* to deliver transcutaneous stimulation, where AM occurs at an area of interest to deliver phasic stimulation at a target region, for instance to stimulate muscle activation. B, We compare suprathreshold electrical stimulation of peripheral nerves using kHz carriers with three different waveforms and four different stimulation conditions: TIS, with 4 electrodes and AMF generated by interference in the tissue; AM sine, with 2-electrodes and AMF given by premodulation; Sine burst, where the kHz stimulus is turned on/off; and finally unmodulated sine. C, The peripheral nerve stimulation models tested in this work. First an insect model: stimulation of the N5 nerve in *Locusta migratoria*, giving evoked movement of the leg as the biomarker. Second, the median nerve in the human forearm. Evoked motor activity was tested by stimulating the median nerve branch to the *flexor digitorum superficialis* muscle, and evoked sensory response was tested by stimulating the median nerve at the carpal tunnel.

### Sinusoidal kHz stimulation and the strength-frequency relation

In scanning both older (ICS) and newer literature on TIS, the assumption that the kHz carrier frequencies (i.e. f1 and f2) do not elicit stimulation themselves has been rationalized in various ways. Often it is assumed that the effect of opposite polarity sinus phases cancels out, since the frequency exceeds that of typical firing rate of neurons. Another rationale for this assumption is based on the impedance/frequency characteristics of neural tissues. Due to the capacitive properties of cell membranes, higher frequency currents pass through cell membranes with lower loss, that is they will induce a lower voltage drop over the cell membrane and thus the membrane experiences lower effective electric field as carrier frequency is increased27. This argument was invoked by Grossman et al. as the “low-pass filtering” explanation14. Another aspect of the “impedance/frequency argument” can be seen in the ICS literature: The concept of higher frequencies encountering lower loss has been used to claim that high-frequency carriers can achieve more effective penetration than conventional pulsed current stimulation, allowing TIS to stimulate deeper targets than conventional (low-frequency) forms of transcutaneous stimulation9,28. While often taken as a given in papers, it has been pointed out that this “depth” argument is erroneous since the phase length of typical biphasic pulsed current stimulation is not different from kHz sine waves (hundreds of microseconds) 11,29. Regardless of the rationale, TIS/ICS studies often assume that the carrier sine itself is not the origin of the stimulation effect. Neural activity preferentially occurs at frequencies < 100 Hz, thus in TIS experiments it is expected that effective stimulation is produced at the AMF which is also in this range < 100 Hz. The central experimental goal of our paper is to test how much stimulation thresholds differ between the amplitude-modulated kHz, and the unmodulated carrier. The key method we rely on is measuring strength-frequency (s-f) dependence. This is the frequency-domain analog to the well-known strength-duration (s-d) dependence of neural stimulation. To the best of our knowledge, a comparative study of s-f dependance in the context of TIS has not been reported before. Historically, s-f threshold characteristics were measured for sinusoidal stimuli into the kHz region30. Sinusoidal frequencies in the kHz range can indeed lead to stimulation, and the associated effect of nerve conduction block (kHz frequency nerve block)31–36. The fact that neurons cannot fire at higher frequencies does not mean they are not electrically stimulated by higher frequencies. Experiments in the 1930s-1960s found that kHz frequencies 0.5-50 kHz would lead to evoked action potentials, and that threshold current required for stimulation increased with frequency30,37–39. Electrodes in direct contact with the nerve can even produce suprathreshold stimulation with sinusoidal frequencies up to 1 MHz40. The s-f curve in the kHz range has been confirmed in different excitable tissues and cell types41. Over the frequency range of sinusoidal stimuli, it is apparent that the s-f relation (Figure 2A) is governed by two exponential functions describing increasing thresholds at very low frequencies (< 40 Hz, the so-called accommodation region), and at high frequencies (around 100 kHz and higher), producing a characteristic U-sharped curve42,43. This sine stimulation U-curve was first reported by Hill in 193644 and Katz30 and described by the Hill equation. The empirical equation for the sine U-curve (plotted in Figure 2A) was later refined by Reilly and colleagues43. The reason why a symmetric sine wave leads to net stimulation is because the depolarizing effect of the first phase is not fully counteracted by the subsequent opposite-polarity phase, due to nonlinear ionic conductivity across excitable cell membranes35,41. This effect can be illustrated by using established computational methods for model neurons, as first pointed out by Mirzakhalili et al.17. Figure 2B shows the calculated asymmetric membrane voltage change in response to a symmetric 1 kHz sinusoidal stimulus (i.e. rectification effect). The sine stimulation leads to net depolarization of the cell. This is a result of asymmetric conduction behavior of sodium and potassium channels. Sodium channels respond faster and allow more sodium into the cell before inactivating, while the potassium channels are slower and less conductive, conducting relatively less potassium out of the cell. This results in more sodium in than potassium out, causing net depolarization with each successive sinus period45. This causes the related effect of summation of subthreshold depolarizations, since at kHz frequencies successive half-cycles arrive before the neuron can restore resting membrane potential via leak currents and other mechanisms. This kind of summation effect for kHz sine stimulation was first described by Gildemeister37, to explain his observations that stimulation efficacy of kHz bursts was strongly dependent on burst duration (aka number of repetitions of the sinusoidal cycle). Using the model neuron, one can observe in practice how subthreshold and suprathreshold stimulation with unmodulated kHz versus TIS looks like (Figure 2C). An unmodulated subthreshold kHz stimulus causes net depolarization and holds that depolarization in a way analogous to DC stimulation (blue trace in Figure 2c). Gildemeister observed this effect in the 1940s and conceptualized a symmetric kHz signal as having the same net effect as applying a DC stimulus at each electrode, calling it “ambipolar stimulation”38,39 since each electrode causes the same net depolarization (there is no “anode” or “cathode” with a kHz sine). He observed that cells below both electrodes were depolarized to the same level, as if both were acting like DC cathodes. As the current amplitude of the kHz sine increase, suprathreshold stimulation is reached (green trace in Figure 2C). The summation effect “build up” can be seen in the low-pass filtered membrane potential trace (dotted green line). The potential rises with successive kHz periods until the membrane potential threshold for AP firing is reached. By applying a Δf= 5 Hz frequency offset between kHz sines, that is TIS conditions, a very similar behavior is shown by the model: the kHz summation effect (red trace in Figure 2C) causes threshold to be reached every 200 ms. Importantly, the mechanism of stimulation originates from the summation of subthreshold depolarizations caused by the kHz carrier. The AMF can modulate how often threshold is reached, however the stimulation itself is not driven by any “low-frequency” E-field at the AMF. It is important to recall that in TIS, in a frequency/power spectrum of two kHz sine carriers, there is no energy at low frequencies. Two recent theoretical papers17,46 maintain that the phasic stimulation effects observable in TIS experiments are driven by the rectification of kHz signals, and thus that off-target effects caused by tonic stimulation by the unmodulated carriers are unavoidable. These papers point out the importance of considering the effect of sustained application of kHz carriers on tissues superficial to the region being targeted by TIS especially since tonic kHz stimulation can cause the phenomenon of nerve conduction block31,34,47. However, it has been postulated theoretically17 and reported experimentally25, that due to resonant coupling of a kHz signal modulated by a “preferred” AMF, a given AMF can result in an optimally low stimulation threshold. Thus, it is possible to have a situation where a given unmodulated carrier is subthreshold, while the region of AM gives suprathreshold stimulation at the beat frequency. Therefore, in this paper we set out to experimentally verify the stimulation thresholds for unmodulated kHz stimuli and compare them with AM kHz and AM delivered by TIS (as summarized in Figure 1B,C). We found that TIS shows the same s-f relation as other modulated and unmodulated kHz waveforms, firmly establishing that TIS is a form of kHz stimulation and there is no evidence for invoking another low-frequency demodulation mechanism. We have however measured some differences between apparent threshold of modulated/unmodulated kHz sine, as predicted by theory.

**Figure 2.**
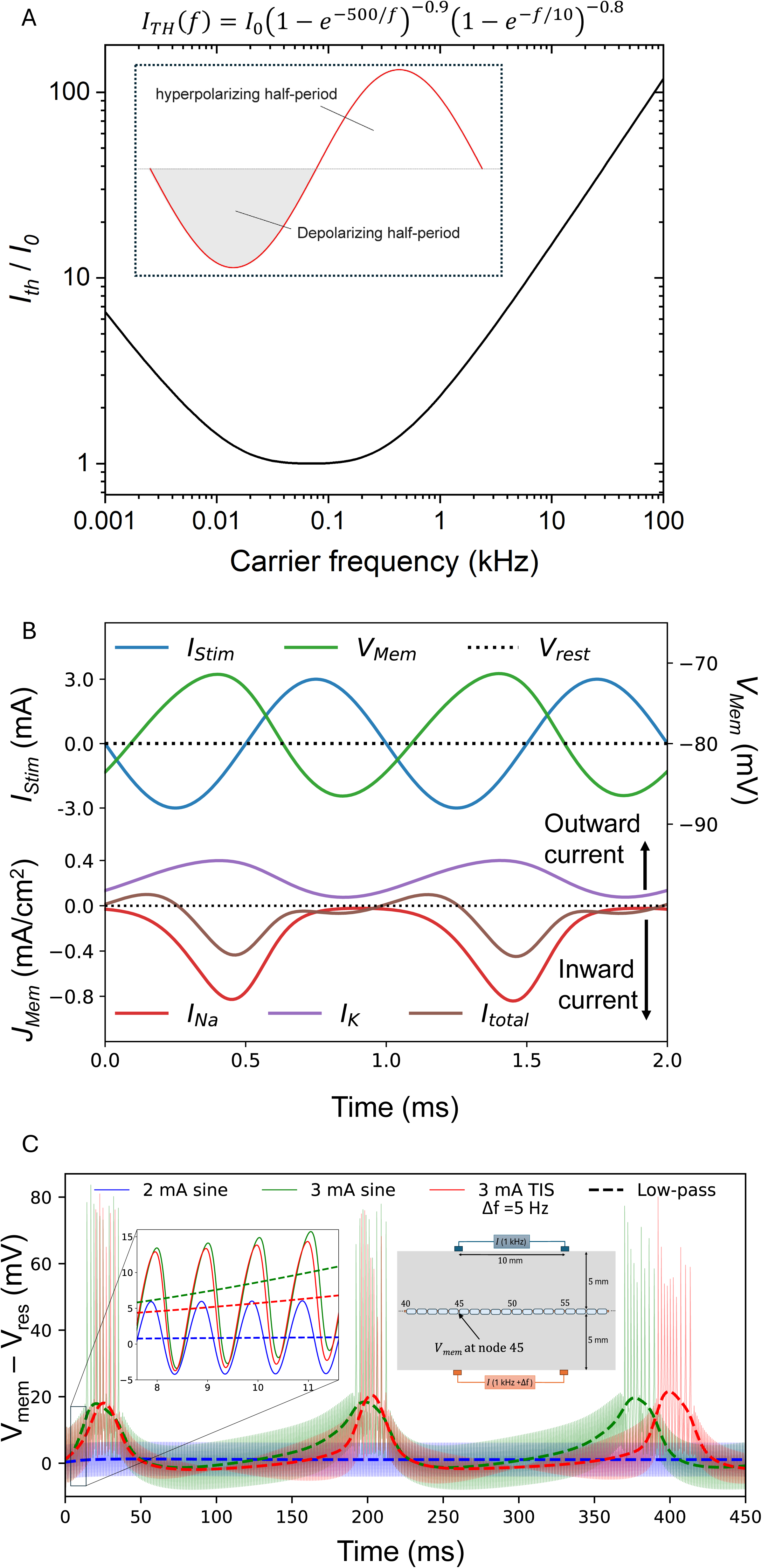
Biophysics of sinusoidal current stimulation. A, The strength-frequency, s-f, relation for the threshold current of sinusoidal stimulation is described by the Reilly equation^43^. The inset schematizes sinusoidal stimulation. It follows that if TIS stimulation is a function of carrier frequency according to this s-f equation, then TIS is driven by the carrier sine itself and not another mechanism. B, According to the Mirzakhalili model^17^, Symmetric kHz stimulation waveforms cause net depolarization of cells, because of membrane rectification. Over one period of the kHz sine, there is more Na^+^ influx than K^+^ outflux, thus leading to net inward current and depolarization. C, 1 kHz sinusoidal stimulation on a mammalian myelinated axon model,^17^ demonstrating the effect of rectification leading to depolarization. The model shows the calculated membrane potential during application of an extracellular stimulating current. Three test cases are shown: unmodulated sine subthreshold stimulation (blue line), unmodulated sine suprathreshold stimulation (green line), and two sines with Δ𝑓 =5 Hz carrier frequency offset (TIS, condition, red line). All solid lines are unfiltered, showing the kHz oscillations, while the dashed traces are low-pass filtered to allow easier visualization of the slower membrane potential changes. Subthreshold stimulation gives a net membrane depolarization analogous to cathodic DC, however when the stimulation current becomes suprathreshold, one can see the summation effect at work as the membrane depolarizes more with each successive sinusoidal period, until threshold is reached and action potentials fire at the preferred resonance frequency of the neuron (5 Hz in this case). TIS, with a 5 Hz AMF, leads to a similar suprathreshold stimulation effect.

## 2. Results

### TIS has the same strength-frequency dependence as 2-electrode kHz stimulation waveforms

To provide rapid prototyping of the TIS protocol versus kHz stimulation, we suggest the use of an invertebrate such as the locust (*Locusta migratoria*). Insects can provide a useful model for noninvasive electrical stimulation, as despite obvious structural and physiological differences to mammals/humans, their cuticle has comparable electrical characteristics to human skin^48^. Moreover, the locust has a relatively simple and well-understood nervous system, including a pair of large symmetric motor/sensory nerves known as the N5 nerves, projecting from the metathoracic ganglion (Figure 3A).^49^ The N5 nerves control movement of the remarkable hind legs of the animal^49,50^. Stimulation of the N5 nerve leads to a highly-reproducible biomarker in the form of evoked leg motion, which can be finely tracked with video analysis^51^. The reason for this reproducible evoked biomarker is that the largest muscle in the leg, the extensor tibiae, responsible for jumping, is controlled by only three neurons. Of the three, the fast extensor tibiae motoneuron (FETi) has an axonal cross section three times larger than any other neuron inside the N5 nerve, thus it is preferentially stimulated, and the downstream actuation of the extensor tibiae muscle is a reliable biomarker. The stimulation of the FETi motor neuron was described by Zurita et al.^51,52^ using a nerve-cuff electrode and tracking protocols for the N5 nerve in the locust, which inspired our method to stimulate the N5 target noninvasively. We applied two electrode pairs aligned with the N5 nerves so that opposite electrode pairs are oriented obliquely with respect to the axon direction. This way, the midway regions between the two electrode pairs where modulation index is maximal will be aligned over the N5. Using TIS in this configuration, we were able to evoke robust actuation of leg motion, using AMFs between 0.1 – 2 Hz. We next compared TIS using this optimized 4-electrode placement for stimulating the N5 nerve, versus a 2-electrode arrangement where the stimulation electrode was placed directly above the N5 nerve (the configurations are compared in Figure 3A). The tested frequency range of the applied kHz signal, or carrier signal, was 500-12500 Hz. As the biomarker for stimulation, we evaluated the motion of the leg, using video to track movement. We measured the carrier frequency-current threshold dependence (s-f curve) for TIS versus the two modulated sine waveforms and the unmodulated sine waveform (Figure 3B). The AMF was set to 1 Hz for all modulated experiments. To ensure an approximately equal time for each trial, the maximum output current was changed from 1 mA, for frequencies below 5 kHz, to 10 mA for frequencies greater than or equal to 5 kHz. In all locust experiments the same ramping procedure was used across waveforms and consisted of starting stimulation at approximately 5 µA and increasing stimulation in increments of 5 µA (<5 kHz) or 50 µA (≥5 kHz) every 0.5 seconds. Once the threshold was reached the current was decreased until the evoked movement was no longer present. At this point the current was increased again to the threshold point and the threshold current recorded. The presentation of a given frequency was randomized for every trial. The modulated waveforms all produced a phasic kicking at 1 Hz, while the sine wave stimulation resulted in a tetanic contraction with the leg remaining in a fully extended position. All four stimulation types demonstrated a very similar s-f dependence (Figure 3B). Statistical analysis was completed using a linear mixed effect model with fixed factors of frequency, waveform and their interaction term, as well as a random factor for ‘participant’. A box-cox transformation was applied to the data (λ = 0.02). This analysis highlighted a significant difference for the main effect of frequency (F(7,92)=940.73, p<0.0001) and waveform (F(2, 92) = 40.70, p < 0.0001) with the interaction term not being significant (p>0.080). A post-hoc pairwise comparison on the significant waveform effect using Tukey HSD highlighted that this effect was driven by a difference in AM sine threshold relative to sine burst (SE = 0.00058, 95% CI [0.00254, 0.00531], t(92) = 6.76, p < 0.0001) and sine (SE = 0.00058, 95% CI [0.00358, 0.00635], t(92) = 8.55, p < 0.0001). To interpret the practical difference in magnitude in threshold values between modulated and unmodulated sine, it is useful to consider the mean threshold ratio ± SEM. We find that unmodulated sine has, on average across frequencies, lower threshold than modulated waveforms: AM/sine = 1.29±0.04; Burst/sine = 1.06±0.02. The 4-electrode TIS was excluded from these statistical comparisons above due to the inherent differences in electrode placements and the fact that ×2 total current is injected into the system, making a direct comparison of threshold value unreliable. The essentially identical qualitative features between all stimulation types, the overlapping s-f relations, and the finding that unmodulated sine has essentially the same threshold for stimulation, indicate that TIS works by delivering kHz stimulation. It is important to underscore that the current threshold values we report are for one carrier only, that is, to achieve the same stimulation effect with TIS versus the 2-electrode modalities, twice as much total current must be used.

**Figure 3.**
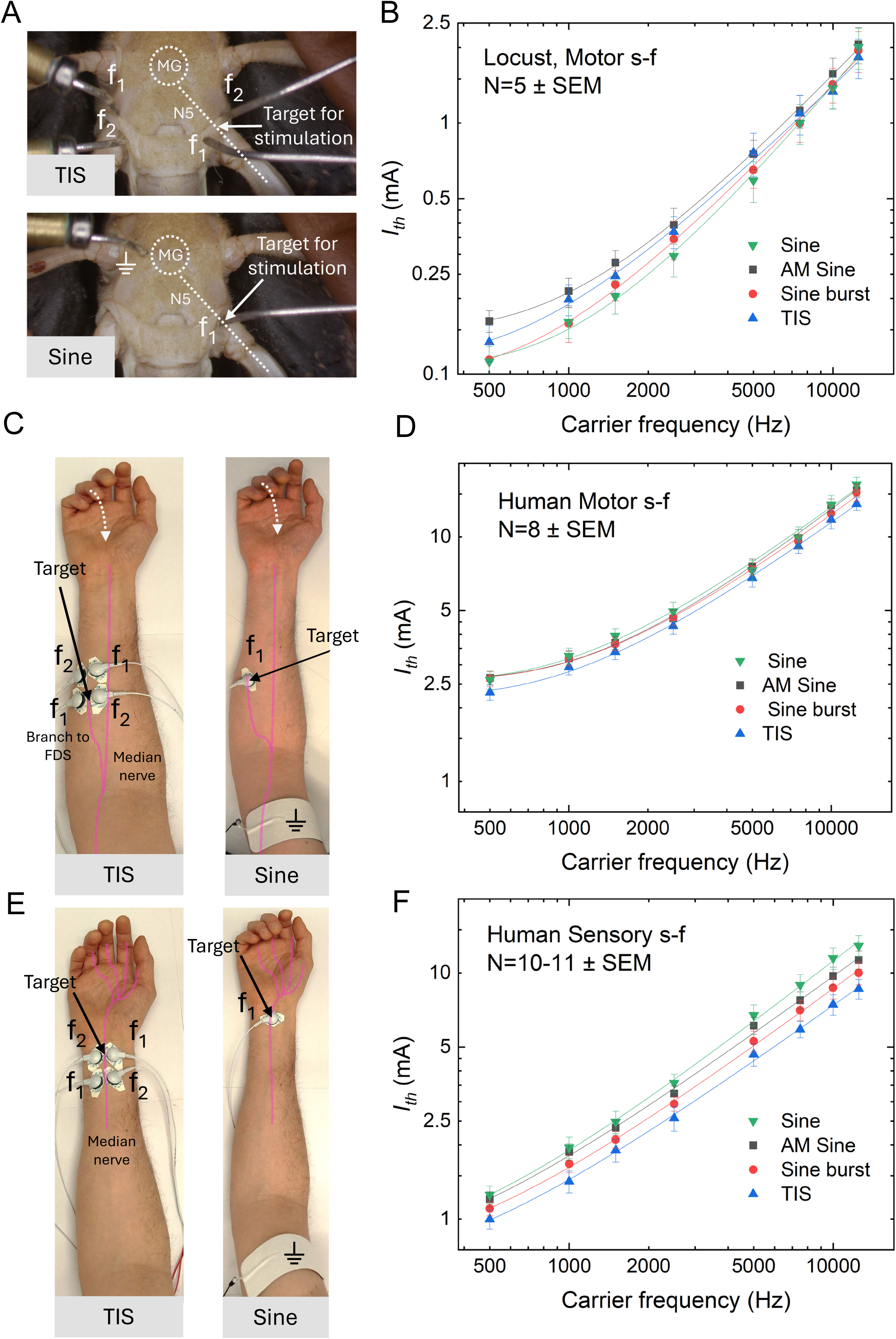
Comparison of strength-frequency (s-f) for 4-electrode TIS versus modulated and unmodulated kHz sinus waveforms in insect and human models. AMF is 1 Hz for all experiments. A, Experimental setup for N5 stimulation, with the two configurations: TIS 4-electrode and “Sine” for all other 2-electrode waveforms. B, Current threshold/frequency s-f dependence for N5 nerve stimulation for the four tested stimulation types. To note: The reported current for TIS corresponds to one carrier only, i.e. the total injected current is ×2 higher. Sine stimulation results in tetanic contraction, while the other waveforms produce phasic motion at 1 Hz. C, Stimulation of the median nerve, comparing TIS with sine waveforms. Stimulation of the median nerve (FDS branch) in the forearm results in reproducible motor response in the form of ring finger flexion a biomarker for stimulation. We arbitrarily define threshold response as the ring finger flexing until it contacts the palm. D, s-f dependence of motor threshold for FDS stimulation for different waveforms with 500-12500 Hz carrier frequency, N=8 participants. All waveforms give phasic motion at 1 Hz, while unmodulated sine produces tetanic contraction. E) Stimulation setup of the median nerve at the carpal tunnel to evoke sensory response in the hand. F) Current threshold/frequency s-f dependence for sensory threshold for the four tested stimulation types. To note: The reported current for TIS corresponds to one carrier only, i.e. the total injected current is ×2 higher. Sine stimulation results in constant sensory response, while the other waveforms produce phasic sensation. All lines between s-f data points are fits of the Reilly equation, R^2^ > 0.99 in all cases.

We next extrapolated the experiment of comparing TIS with direct kHz stimulation to human subjects (Figure 3C-F). As a convenient stimulation target, we chose the median nerve in the forearm. Both afferent and efferent fibers were targeted. Efferent fibers innervating the FDS in the mid forearm were the target for the motor stimulation experiment. To quantify stimulation, flexion and movement of the ring finger was used as a biomarker. Afferent fibers below the carpal tunnel at the wrist were also targeted which evoked a ‘pins and needles’ like sensation in the fingertips and palm (Figure 3C). The threshold, for the efferent fibers, is defined as the current needed to achieve complete flexion and contact of the finger to the palm of the hand (Figure 3E). The threshold, for the afferent fibers, is defined as the onset of ‘pins and needles’ in the tip of the participant’s ring finger. For TIS, the median nerve was optimally targeted by placing opposite stimulation pairs such that the nerve lay along the midline between the electrodes, analogous to the electrode placement used for the locust N5. The location of the TIS electrodes was then used to define the position of the electrodes for two-electrode kHz waveforms, the electrode was placed directly over the median nerve, and stimulation was applied versus a relatively large ground electrode. We performed stimulation using carrier frequencies between 500-12500 Hz, first with AMF of 1 Hz. To ensure an approximately equal time for each trial the incrementation step size was set differently for frequencies below 5 kHz compared to frequencies greater than or equal to 5 kHz. The same ramping procedure was used for each waveform and in both median nerve models. It consisted of increasing stimulation current from 0.25 mA in increments of 0.025 mA (< 5kHz) and 0.25 mA (≥5kHz) every 0.5 seconds until threshold motor activation was reached, then the current was decreased until the evoked movement no longer persisted. At this point the current was again ramped in the same fashion until the threshold was reached; here the threshold current was recorded. The presentation of frequency was randomized for every trial. Statistical analysis was completed using a linear mixed effect model with fixed factors of frequency, waveform and their interaction, as well as a random factor for ‘participant’. A box-cox transformation was applied to the data (λ = 0.5). This analysis highlighted a significant difference for the main effect of frequency ((F(7,158)=643.51, p<0.0001)) with the main effect of waveform ((F(2,158)=2.70, p=0.0700)) and the interaction term (p>0.087) not being significant. Therefore, stimulation thresholds across waveforms and the entire frequency range tested are not statistically significantly different from each other. The average threshold ratios for modulated/unmodulated sine waveforms are equivalent across the frequency range: AM/sine = 0.99±0.02;

Burst/sine = 0.97±0.02. Further the s-f dependance for all stimulation modes was found to be the same (Figure 3D), and the Reilly equation (as shown in Figure 2A) fit to each case. As before with the locust model, qualitatively the movement evoked by each stimulation was the identical. Participants reported that the sensation accompanying AM sine and TIS was essentially similar, with minimal or no skin sensation at the site of stimulation. The Sine burst was accompanied by obvious skin sensation and was subjectively less comfortable than the AM sine and TIS stimulation patterns. These differences in subjective perception could be related to the degree of synchrony that the different modulated waveforms generate^53^, and this a topic of current study and beyond consideration of this manuscript however, and we litigate only the threshold of sensory perception or motor movement. In our afferent median nerve model, an identical statistical approach was taken as for the motor stimulation. The only difference being that a box-cox transformation was applied with λ = 0.19. The other features of the model were the same as stated above. This analysis highlighted a significant difference for the main effect of frequency (F(7,221.01)=1146.55, p<0.0001)), waveform (F(2,221.12)=43.15, p=0.0700)) with the interaction term not being significant (p>0.179). A post-hoc pairwise comparison on the significant waveform effect using Tukey HSD pairwise comparison highlighted that this effect was driven by a difference in all three waveforms pairs, sine burst – AM sine (SE = 0.00537, 95% CI [−0.03876, −0.01340], t(92) = −4.85, p < 0.0001) and sine – AM sine (SE = 0.00520, 95% CI [0.01174, 0.03628], t(92) = 4.62, p < 0.0001) and sine – sine burst (SE = 0.00539, 95% CI [0.03737, 0.006281], t(92) = 9.29, p < 0.0001). These differences equate to ratios of sensory thresholds which favor the modulated waveforms: AM/sine = 0.93±0.02; Burst/sine = 0.84±0.02. This trend is clearly visible in figure 3F and we would maintain that this is intrinsic to using reported subjective perception as a read-out. Simply put, signals that turn on/off in a rapid way provide a sensory contrast that is easy for people to perceive. For sine burst, the sharp onset of the waveform is a clear indicator of when threshold is reached, this effect is dampened slightly due to the smoother amplitude modulation of AM sine and the lack of modulation in the sine condition (save for the slow ramping) makes decerning the exact moment of perceptual onset more difficult. TIS has the average lowest threshold, however we would indicate that this is due to a combination of its modulated nature and the 2× applied current, making it the clearest perceptual experience. Despite this caveat, again, a qualitatively identical s-f dependence is observed across all waveforms further adding to the evidence that the mechanisms underlying TIS and kHz stimulation is identical in afferent nerve fibers – the nerve fiber type with the typically lowest threshold for electrical stimulation. Taken together, these median nerve results evidence that TIS operates mechanistically via kHz stimulation. It is important to note that once again, as with the locust model, the current thresholds are reported with respect to one current source only, thus, to achieve the same stimulation outcome, TIS must use ×2 the current as the two-electrode configurations. Second, it is notable that with unmodulated sine, not only was the s-f relation the same as for the modulated waveforms, but the evoked motor movement was identical too, with the exception that contraction of the finger was followed by sustained tetanic contraction in all participants. In practical terms for 4-electrode TIS, this means that the current threshold required for phasic stimulation is the same as tonic stimulation. This question of phasic/tonic effects will be elaborated further in upcoming sections of this article.

### Modulated and unmodulated sine frequencies in the range 10 - 100 kHz share a common s-f relation

To further test the hypothesis that TIS is in principle a type of kHz stimulation and no carrier-independent envelope demodulation mechanism is occurring it is interesting to extrapolate the s-f curve to even higher frequencies. Using a high-frequency voltage source we attempted to apply modulated (AMF = 1 Hz) and unmodulated carriers in the range above 12.5 kHz and even up to 1 MHz. We found reproducible stimulation up to 100 kHz (Figure 4). The Reilly s-f relation continues to hold up to this frequency, and we found no significant difference between AM and unmodulated sine. This was determined using the same statistical model as described above for the previous locust data with a main effect of frequency being significant ((F(7,57.18) = 216.8, p<0.0000)) and the main effect of waveform not being significant ((F(1,57.01) = 1.53, p<0.2214)). We note that it was possible to observe stimulation at even higher frequencies, however results were unreliable and obviously there were effects coming from Joule heating (including visible damage to insect tissue) due to the relatively high currents required to achieve threshold. Therefore, we report only the safe window we found up to 100 kHz. The fading of neural response at higher frequencies, and transition into a thermal dissipation regime, is reported in several studies of kHz neurostimulation^36,40,42^. The fact that the s-f relation of modulated and unmodulated carriers remains overlapping is evidence pointing towards the conclusion that, as with “conventional” kHz frequencies, TIS acts via carrier signal rectification and there is not another carrier-independent mechanism of low-frequency envelope demodulation emerging at increasing carrier frequencies.

**Figure 4.**
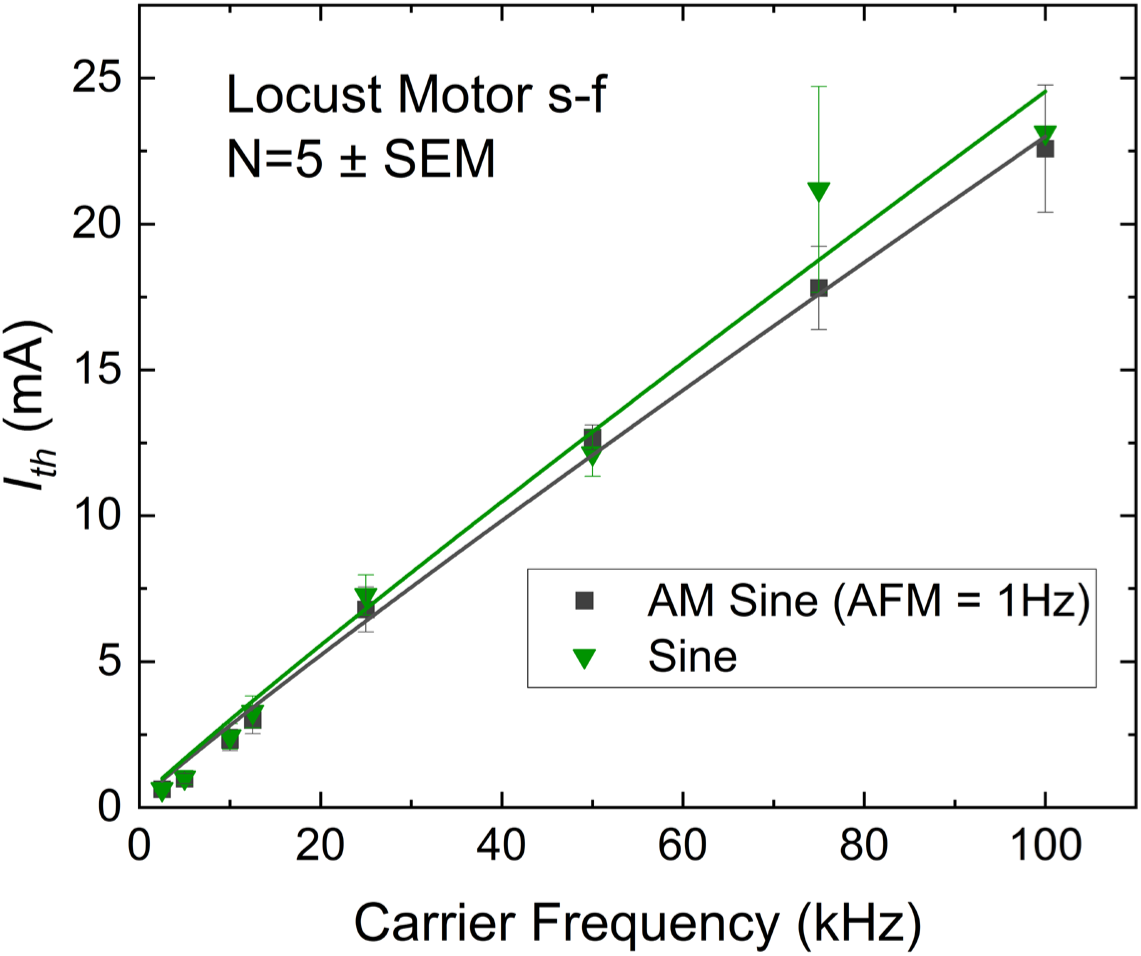
Strength-frequency relation for AM sine versus unmodulated sine for frequencies up to 100 kHz. The locust N5 nerve can be reliably stimulated with carrier frequency up to 100 kHz, in a tonic or phasic way using sine or AM-sine, respectively. Frequencies higher than 100 kHz result in excessive heating and unreliable results.

### Varying AMF has limited impact on the suprathreshold stimulation current threshold

From assessing the s-f relation of TIS and different modulated/unmodulated kHz waveforms, it would appear that threshold current varies strongly with carrier frequency, but at least at AMF=1 Hz, there are no meaningful threshold differences caused by amplitude modulation itself. We next performed a series of experiments to interrogate the effect of AMF in locusts (looking at low AMF below 1 Hz), and in the human forearm models, in a larger range between AMF = 0.03 Hz and AMF = 50 Hz. In locusts, we carried AMF down from 1 Hz to 0 Hz, while keeping the carrier frequency fixed at 2500 Hz. In Figure 5A, we plot the time of initiation of evoked kick and the associated threshold current as a function of the AMF envelope. The threshold does not change significantly with AMF, and the kick is initiated once a common threshold value is reached. During the experiment, we monitored the deflection angle, θ, of the femoral-tibial joint, defining Δθ as the difference between the unstimulated resting angle and the maximum deflection angle evoked by stimulation (Figure 5B). Stimulation at the minimum threshold current at AMF = 1 Hz gives a Δθ of 5°, and a phasic movement with a period of 1 second. As Δ*f* is lowered, the value of Δθ increases. By AMF = 0.1 Hz, the Δθ has increased to 34°, with a phasic movement every 10 seconds. What is apparent is that the lower the beat frequency is, the longer the N5 nerve is subject to suprathreshold kHz stimulation, thus causing the *extensor tibiae* muscle contraction to be sustained for a longer period, giving a larger Δθ. The time over threshold, plotted on the right y-axis, corresponds to the deflection angle. This point is made more evident by decreasing the AMF to 0 Hz, that is, stimulation with unmodulated 2500 Hz. The leg then deflects to a maximum Δθ of 57° and remains at this position in a tetanic contraction. This experiment aligns with the original observation of Gildemeister^37^, that the stimulation effect is proportional to the duration of application of kHz pulses, known as the temporal summation effect. Moreover, the effect of having the stimulation set to the threshold for suprathreshold kHz stimulation and the transition between tonic and phasic stimulation can be neatly demonstrated as shown in Figure 5C: The lowest threshold for phasic movement at 1 Hz is set but using a 1 Hz modulated sine stimulation. The leg then produces a Δθ of 5° movement with a frequency of 1 Hz. Next, while keeping the current amplitude the same, the modulation is turned off, tonic stimulation will result, and the leg extends in tetanic contraction. The phasic/tonic problem will be discussed in more detail in the next section.

**Figure 5.**
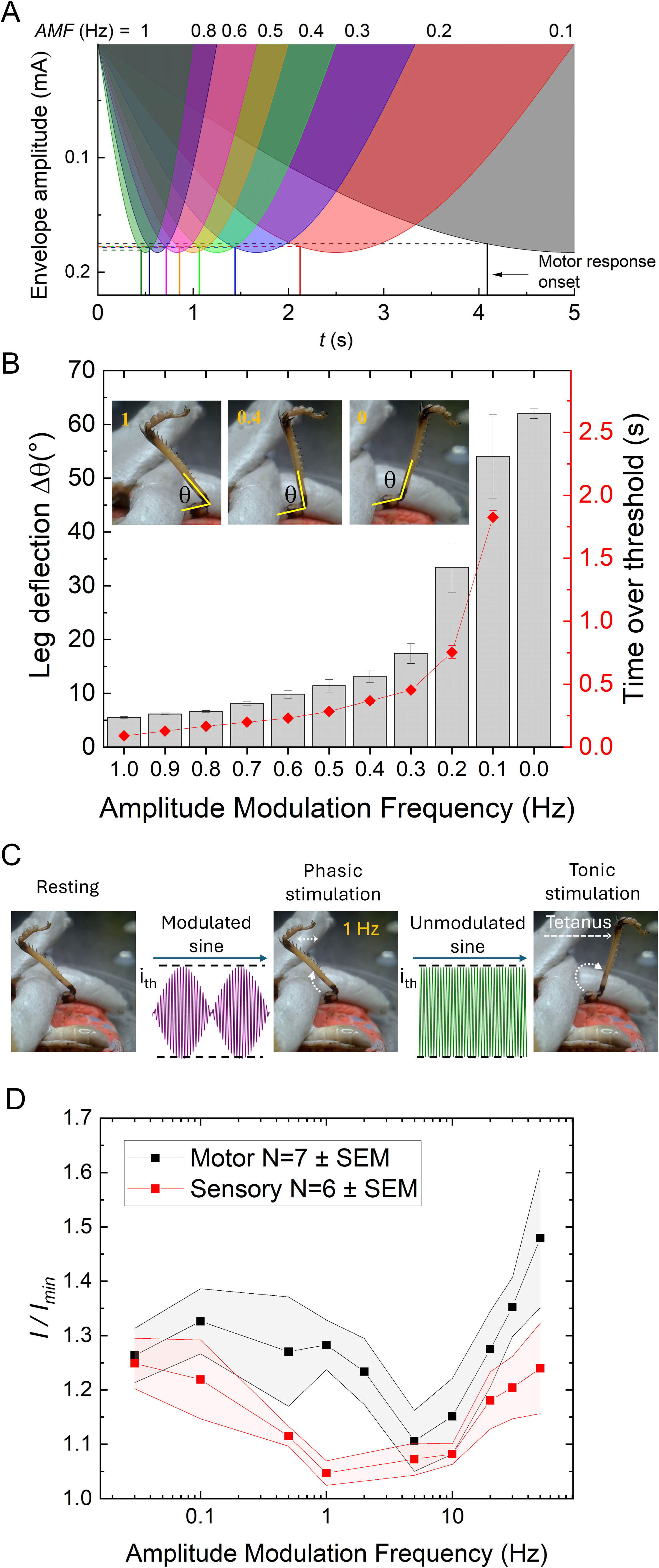
Stimulation thresholds as a function of amplitude modulation frequency, AMF. A, Locust N5 nerve stimulation threshold current and timepoint of evoked movement as a function of AMF, plotted on top of the AM envelope of the kHz stimulation (carrier frequency = 2.5 kHz). In this range of AMF, the threshold does not vary with AMF, the kick is evoked once a constant current value is reached, regardless of the amplitude ramping conditions. B, Leg deflection angle, Δ*θ*, caused by *extensor tibiae* actuation, as a function of AFM. The deflection angle correlates to the length of time that the kHz stimulation is over threshold, with unmodulated kHz giving the greatest deflection angle change. Mean value ± SEM for each AMF, N=9-23 experiments. C, This experiment involves applying an at-threshold stimulus at AMF = 1 Hz, and then turning off the modulation, while maintaining the current setpoint. This causes tetanic contractions. Turning the modulation on and off will shift the stimulation result from phasic movement to tetanic contraction. This test shows that threshold current for the phasic stimulation and the tonic stimulation are equal. D, Effect of AMF on the motor and sensory thresholds in the median nerve stimulation for a fixed stimulation frequency of 3000 Hz. The characteristic U-shaped response predicted by theory is demonstrated, with statistically significant increases in threshold at low and high AMF, relative to the optimum AMF = 5 Hz.

The model of nonlinear neuronal membrane conduction properties which result in rectification of sinusoidal signals also predicts that this rectifier circuit responds to an optimum AMF, aka the “preferred frequency” of the neuron.^17^ That is, a TIS frequency that gives the lowest activation current threshold.^17^ For myelinated axons, the model predicts a U-shape of current threshold versus AMF, where an optimal minimum of current threshold should exist at a given AMF, and very low or very high AMF will lead to higher current thresholds. This optimal AMF is predicted to be in the range of 5-40 Hz. Experimental data, however, suggest that this effect may be weak in peripheral nerve stimulation. In his 1939 paper, Katz reported no threshold difference between rapid and slow amplitude ramps on kHz stimulation of the frog sciatic nerve.^30^ In a study of sensor, motor, and pain thresholds using ICS, Palmer et al., found lack of dependence of thresholds on AMF, although this study used a window of frequencies of 5-100 Hz.^54^ Budde et al.^25^ using an implanted cuff electrode on the rat sciatic nerve, did demonstrate a U-curve dependence with AM-kHz stimulation (AMF range 1-1000 Hz) of the sciatic nerve, with dependence roughly ±10 – 20% lower thresholds for AMF = 10-50 Hz than for very slow or fast AM. We performed experiments on the human median nerve to test AMF dependence from very low frequencies approaching unmodulated sine waves (0.03 Hz) up to 50 Hz. We selectively stimulated sensory fibers at the location of the carpal tunnel to establish sensory threshold (setup from Figure 3E), and motor threshold by stimulating the median nerve branch to the FDS at the upper forearm (setup in Figure 3C). In both cases, we did observe a shallow U-shaped dependence on current threshold with an apparent optimal beat frequency of AMF = 5 Hz in the motor nerves and AMF = 1 Hz in the afferent nerves (Figure 5D), where thresholds were lower by roughly 20% compared to low or high AMF. Participants were presented with randomized AMF signals, to eliminate any effects of ascending or descending AMF presentation. Due to variation of absolute threshold values between participants, the data are normalized *i/imin* for each participant, then these values are averaged and evaluated. Not every participant had the *imin* value at the same AMF, thus the averaged normalized curves do not intersect exactly 1 at any point. Statistical analysis using a linear mixed effect model was employed to determine if the apparent observed decrease in threshold was statistically significant. A fixed factor of modulation frequency and a random factor of participant was set. For the sensory model it was found that AMF = 1 Hz was significantly different to frequencies, 0.03 Hz (p < 0.0031), 0.1 Hz (p < 0.0121)), 20 Hz (p < 0.0405), 30 Hz (p < 0.018935) and 50 Hz (p < 0.0058). AMF = 5 Hz was found to be significantly different to frequencies, 50 Hz (p < 0.1590), 30 Hz (p < 0.0471), 0.1 Hz (p < 0.0312) and 0.03 Hz (p < 0.0089). This aligns with the observed pattern in figure 5D and its less ‘sharp’ dip compared to the motor data. An identical statistical model was used to determine significance levels in the motor nerve model. For AMF = 5 Hz the following frequencies were found to be significantly different, 50 Hz (p < 0.0004), 30 Hz (p < 0.0074), 1 Hz (p < 0.0442) and 0.1 (p<0.0157). For AMF = 10 Hz the following frequencies were found to be significantly different, 30 Hz (p< 0.0282) and 50 Hz (p< 0.0019). Overall, the “U-curve” can be considered borderline significant.

### TIS intrinsically gives bimodal stimulation with tonic and phasic regions, while two-electrode kHz stimulation is monomodal

In TIS stimulation, the kHz carriers are unmodulated, and amplitude modulation arises due to interference in the tissue. As a general rule, maximum depth of AM will occur at midpoints between electrode pairs. This means that regions of tissue superficial to the point of maximum AM will be exposed to kHz carrier stimulation with a lower index of modulation. Moreover, since electric field amplitude will fall with distance from the stimulation electrode, these areas could be exposed to relatively high magnitudes unmodulated kHz carrier. This will thus cause regions of phasic simulation (AM region where interference is optimal), and regions of tonic stimulation (regions closer to the stimulation electrodes). We refer to this coexistence of tonic and phasic regions as *bimodal stimulation*. This intrinsic issue with TIS has been pointed out by other researchers.^17,25,46^ The phasic/tonic effect can easily be demonstrated in the FDS motor stimulation experiment in the forearm (Figure 6A, supplementary video 1). An optimal placement of the 4 TIS electrodes, producing maximum AM at midway points between electrodes, will cause phasic muscle activation. While keeping the stimulation current conditions the same, moving one of the 4-electrodes by 1 cm, as shown in Figure 6A, will cause a shift to tonic stimulation and tetanic contraction of the muscle. We use finite element method in a forearm model to explore and visualize the implications of our experimental findings that the stimulation thresholds do not meaningfully differ for modulated and unmodulated stimulation (Figure 6B). We perform frequency domain simulations and calculate the relevant activating functions in the nerve direction 𝐴1 = |𝑑^2^𝑉1/𝑑𝑥^2^| and 𝐴2 = |𝑑^2^𝑉2/𝑑𝑥^2^|, where 𝑉1 and 𝑉2 are electric potential amplitudes generated by stimulation currents 𝐼_1_ and 𝐼_2_ respectively. ^55^ The simulation details can be found in the Computational modeling section. We plot the maximum amplitude 𝐴_MAX_= 𝐴1 + 𝐴2 in the horizontal cross section at the nerve depth which is assumed to be 1 cm below the skin surface (Figure 6C). We use our experimental finding of a unique stimulation threshold 𝐴𝑇 to quantify and visualize the stimulation response mode (tonic, phasic, or subthreshold) in the various regions in the tissue. For this purpose, we use time over threshold (TOT) to mean the fraction of each modulation period that the nerve is suprathreshold, mathematically defined as:

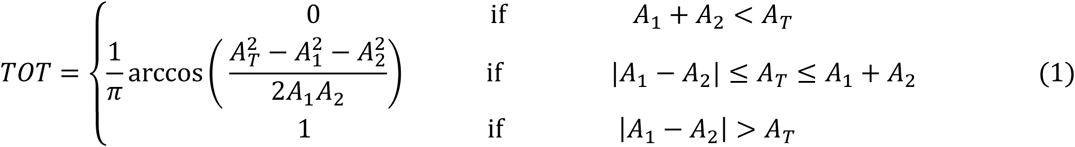

**Figure 6.**
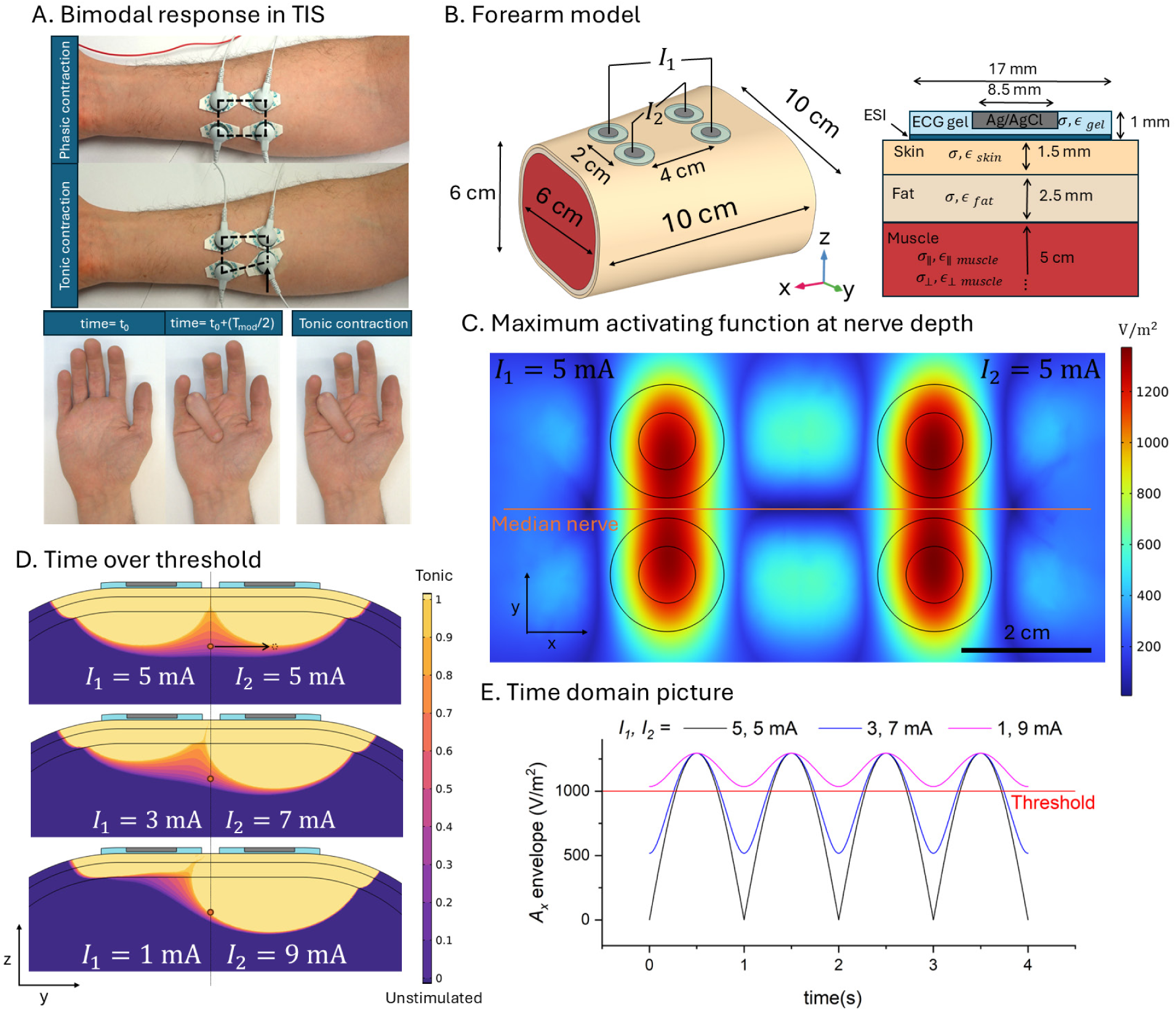
Simulated tonic versus phasic stimulation regions in the model human forearm. A, Demonstration of the bimodal phasic/tonic response in TIS configuration. Moving one of the 4-electrodes directly over the nerve will result in tonic stimulation and tetanic contraction of the muscle (see also supplementary video 1). B, Geometry and the 3-layer tissue model used for simulation. Electrode-skin interface (ESI) is modeled by a distributed impedance consisting of constant phase element in parallel to resistance. C, Horizontal cross section showing the activating function maximum at the nerve depth, assumed to be 1 cm from the skin surface. D, Vertical cross section showing the time over threshold (TOT) for different stimulation current ratios (current steering), and the spatial distribution of tonic (1) and phasic (0.5) regions. Moving the electrodes directly over the nerve shifts the response from phasic to tonic, as observed experimentally. E, As the current ratios change, the nerve is exposed to different degrees of modulation, with the ratio 1, 9 mA causing the nerve to be stimulated over threshold in a tonic way.

We obtained the value for the stimulation threshold 𝐴𝑇 in the FEM simulation by using the experimental current threshold measurements from Figure 3D. For 2.5 kHz carrier frequency calculated stimulation threshold 𝐴𝑇 is 1000 V/m^2^. TOT = 0 represents an unstimulated region where stimulation is never over the threshold. TOT = 1 is the tonic region where stimulation is always over the threshold while values between 0 and 1 define the phasic response region where the stimulation periodically changes from subthreshold to suprathreshold. TOT for different values of stimulation currents 𝐼_1_and 𝐼_2_ is shown in Figure 6D. It reveals regions around the electrodes which have a high unmodulated activating function (TOT=1). Nervous tissue in these regions would be subject to tonic stimulation. In between the electrodes, there are regions of maximum modulation index of the activating function. Here, phasic stimulation will be observed at the beat frequency Δ*f*. These calculations align with what is observed experimentally, as shown in Figure 6A. Time domain representation of activating function envelope at the nerve location for current ratios shown in Figure 6D are shown in Figure 6E.

Since we have observed the essential equivalence of stimulation effects with TIS and AM-kHz, it can be useful to apply finite element modeling to visualize key similarities and differences between the two stimulation approaches (Figure 7). We make a simple geometry to illustrate some general points: Assuming a uniformly-conductive medium with circular symmetry, one can see that performing 4-electrode TIS or 2-electrode AM sine will lead to equivalent stimulation in the center of the model: a phasic stimulus with the same amplitude and maximum modulation index. The critical difference is that TIS will require 4 electrodes and thus twice as much total current injection, and will create regions of relatively high-intensity tonic stimulation below the respective electrodes. This results in bimodal stimulation – some areas are phasic, some are tonic, and areas in between will have a gradient in amplitude modulation index. In 2-electrode sine stimulation, all regions are exposed to the same phasic stimulation with identical modulation index. The attenuation of the E-field with distance will be the same in both TIS and 2-electrode sine configurations. This we refer to as monomodal stimulation. It should be noted that in practical experiments, like we have done in Figure 3, one of the two electrodes in AM sine can be much larger than the primary stimulation electrode, thus limiting unwanted stimulation at that electrode site by decreasing current density. In experimental situations where one wants to avoid tonic stimulation effects, a 2-electrode pre-modulated configuration may be preferred over the 4-electrode TIS.

**Figure 7.**
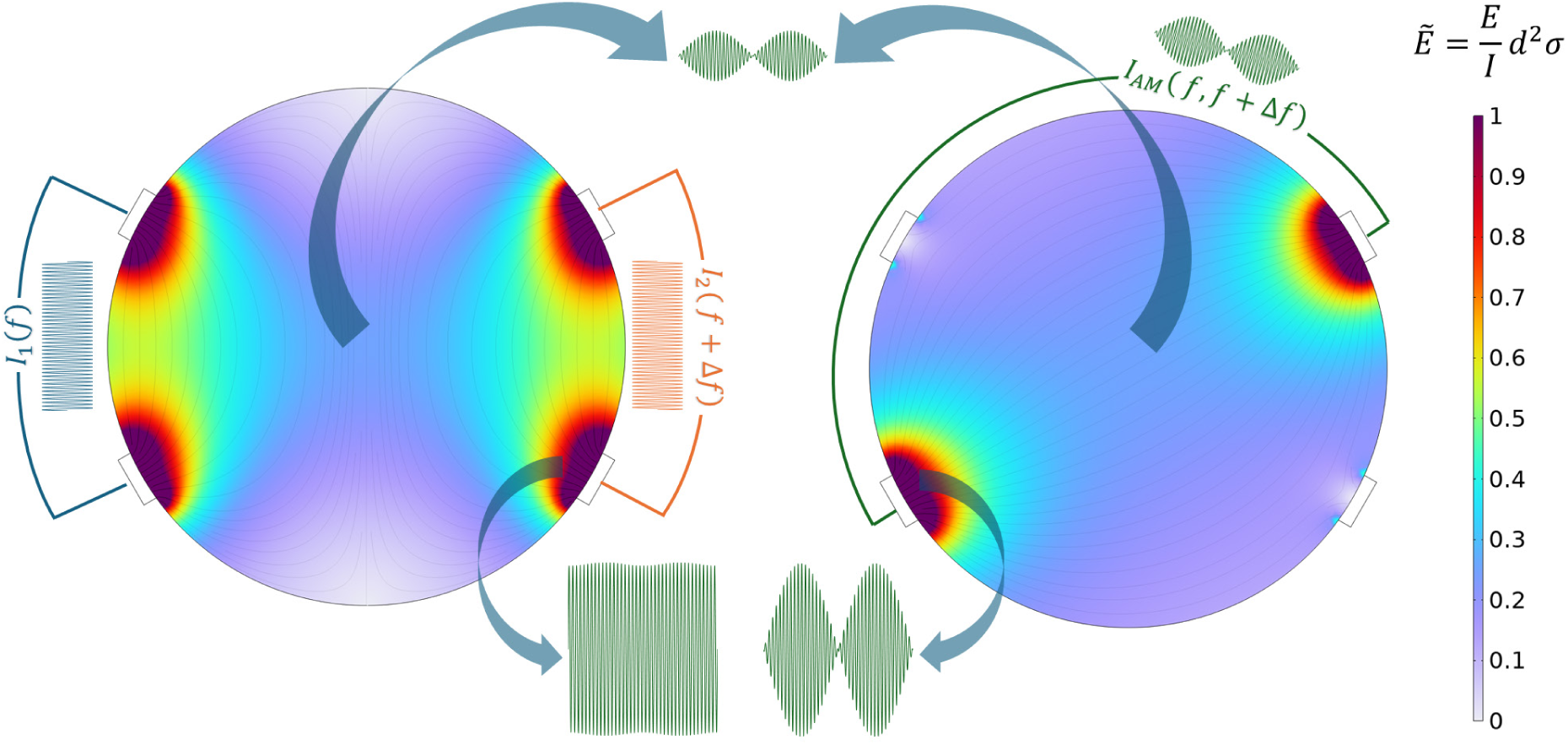
**Generalized calculation of the scaled electric field Ē, comparing TIS with a 2-electrode AM-kHz stimulation** in a circular medium of diameter *d*, conductivity σ, stimulation current *I* and electric field amplitude *E*. The value of the scaled electric field Ē is only a function of the geometry and is independent on the chosen values of *I*, *σ,* or spatial scale *d*. This picture explains why TIS will always result in higher amplitudes of unmodulated carriers superficial to the region of maximum amplitude modulation where phasic stimulation is delivered. Both stimulation methods lead to the same amplitude of phasic stimulation in the middle of the medium.

## 3. Discussion and conclusions

The TIS (*aka* ICS) method has been proposed as a noninvasive electrical stimulation technique which can yield focused stimulation at a target region at-depth. In recent papers describing the use of this method, it is assumed at the outset that the kHz frequencies used as carriers are not able to elicit stimulation on their own, at least not at the amplitudes being used in the experiments.^14,19,56,57^ The experiments we are reporting, as well as recently published *in vitro* studies,^58,59^ indicate that this assumption is not correct. We find no evidence for selective extraction of the envelope AMF, but rather that the biophysics of stimulation originates from the sinusoidal carrier itself. Our work, corroborated by these *in vitro* studies, shows that the threshold for stimulation by kHz sine waves varies little, if at all, with amplitude modulation. The hypothesis that unmodulated and modulated kHz likely have similar thresholds can be anticipated from older literature, starting as early as Bernhard Katz in 1939 who reported no effect of ramp rate on threshold for kHz stimuli on a frog sciatic nerve.^30^ Symmetric periodic signals in the kHz range will lead to net depolarization of neurons and can in fact stimulate. Modern scientists may overlook this due to the age of these studies (most of the work was published between the 1930s and 1970s), and the fact that the foundational papers by Gildemeister^37^ and later Bromm^39,60^ and Wyss^38^ are not available in English and were rarely cited in later English-language literature. As shown in these works, the time of application of the signal matters, due to the principle known as “temporal summation”, the “summation effect”, or “Gildemeister effect”, where multiple successive subthreshold stimuli result in net suprathreshold stimulation.^39,61^ We find, in three different peripheral nerve models (in an insect model and in humans), that direct neurostimulation is driven directly by the kHz stimulus itself, and that the biophysics behind stimulation in TIS versus a single sine wave appear to be the same, *i.e.* the characteristic s-f curve which is established in the literature. While the notion that kHz carriers cannot lead to net stimulation is not correct, it is possible when using optimal AMF (∼ 5 Hz) to exploit the resonance effect to have a situation where the threshold current for the modulated signal is up to 20% lower than for the unmodulated carrier. However, the question is how practically relevant this drop in effective threshold is. TIS for brain stimulation attracted interest because of the promise of selectively targeting deep neural structures, without stimulating superficial structures. If an optimal AMF is chosen, would the resulting threshold drop counteract the effect of E-field decay with distance from the electrode? It would seem that stimulating a deeper target with TIS without exposing the interlaying tissue to higher amplitudes of tonic stimulation is an unlikely prospect and minimally must be carefully considered during the interpretation of the results. A recent *in vitro* study has shown that although the stimulation threshold for modulated vs. unmodulated sine may be the same for a single cell, due to inhibitory network effects TIS may yield differential stimulation of excitatory cells due to input from inhibitory ones.^59^ There is continuing debate on the safety parameters for transcranial TIS^62^ as well as the determination of a “sham” condition for TIS experiments. Based on the potential of the unmodulated sine carrier to stimulate neural tissue, it is arguable if this can be considered a “sham” condition versus the experimental condition of interfering currents and AM.

Our experimental design was motivated to study relatively simple biomarkers, and we focused on suprathreshold stimulation phenomena only. One limitation of our study is that it does not shed light on what can be going on at the subthreshold level. From theoretical calculations (as shown in Figure 2C), one expects that the ion channel rectification phenomenon will be occurring in the same way for subthreshold stimuli, and the cell will net depolarize with similar dynamics as one sees for suprathreshold stimulation. However, our work does not exclude the fact that other, parallel mechanisms related to AMF may in fact be operating in a meaningful way at the subthreshold level. This paper emerged in an effort to address two, disparate communities: the functional electric stimulation community, which has used ICS (this term dominates over TIS) for decades. ICS is standard in clinical practice, and claims about better depth penetration and assumptions about unmodulated sine not stimulating prevail in much of this literature. We believe our results confront these issues, related to suprathreshold stimulation, in an unequivocal way. On the other hand, the TIS community working on brain stimulation is more and more focused on subthreshold stimulation in humans, where other complex phenomena may be at work. There are principled reasons to expect neural processing of amplitude-modulated inputs, for instance the well-known example of auditory steady-state response^63^. In this way our results may be considered limited, however at minimal they should serve to confront the assumption that high-frequency carriers could be electrophysiological inert.

What is remarkable compared to other modulated kHz stimulation is that TIS can deliver bimodal stimulation. That is, there will be tonic and phasic regions, with gradients of amplitude modulation depth in between. An advantage of TIS over a 2-electrode kHz stimulation configuration is the opportunity to use current ratios to steer the position of AM hotspots, and/or exploiting the 180-degree phase shift that will exist between hotspot regions perpendicular to electrode pairs. These features allow interesting degrees of control of phasic/tonic stimulation. These advantages have been nicely documented by Budde et al.^25^ However, this has to be considered in the context of the intervening tissue, which is under such conditions necessarily exposed higher amplitudes of kHz signal with a lower index of modulation. If the target is a peripheral nerve below the skin, with no intervening nervous structures, the TIS method could be advantageous for noninvasive neurostimulation.

In conclusion, while TIS can be a powerful tool to direct stimulation to a region of interest, it is not simply a surrogate for delivering low-frequency stimulation. In the literature, there has been controversy as to the mechanism of how TIS works. Many studies have oversimplified the picture of TIS, maintaining that the high-frequency carriers are inert. Our experiments provide evidence that this picture is incorrect. We conclude that the stimulation observed when using TIS can be explained by conventional kHz stimulation models, and encourage revisiting older literature on the topic. This mechanistic basis must be considered in planning safe and effective TIS applications. Moreover, in many cases, we would argue that a premodulated kHz stimulation arrangement may provide some key benefits of TIS but in a monomodal way, eliminating tonic stimulation effects.

## 4. Experimental section

### Ethics

Experiments on locusts, as invertebrates, do not fall under legislation in the Czech Republic as animal experiments, therefore no special ethical permission is required (Act No. 246/1992 Sb.). Experiments on human healthy volunteers were performed in accordance with approval from the CEITEC Ethical committee for research on human subjects. Stimulation is delivered using the Digitimer DS5 device which is clinically-approved in the EU.

### Electrical stimulation hardware

The kHz stimulation experiments were conducted using the Digitimer DS4 (for locust experiments), and the Digitimer DS5 (for experiments on human participants). For high-frequency experiments in the range 10-100 kHz, we used an A-301HS amplifier from A.A. Lab systems Ltd. Stimulation electrodes for locust experiments were Ag wires (0.6 mm diameter) with chloridated tips (made via immersion in commercial chlorine bleach solution). For human transcutaneous stimulation, the primary stimulation electrodes always consisted of commercial ECG Ag/AgCl gel-assisted adhesive electrodes. For 2-electrode modulated kHz experiments, we used larger return electrodes consisting of gel-assisted carbon TENS pads. Stimulation waveforms are generated by a 2-channel function generator (Keysight Technologies EDU33212A). Synchronization of clock-circuit for each of the channels should be performed via the internal software prior to each experimental session. For premodulation of a sine wave stimulus (AM-sine), two protocols were used: either internal AM modulation provided on one channel of the function generator (AM by multiplication), or by having each channel output a carrier and then mixing the carriers in a BNC T-spitter, which was then fed into the DS5 or DS4 (AM by addition, essentially mimicking the AM created in TIS). While we noted no differences in results from the two methods of AM generation, most experiments were carried out by the carrier mixing for convenience and to more faithfully compare with TIS conditions. Both current simulators (DS4 and DS5) suffer from current amplitude roll-off with increasing frequency > 1 kHz, as can be found in the documentation for each device. This effect was described in detail by FallahRad et al.^33^ To correct for this roll-off, the actual output stimulation current was always monitored in all experiments by registering voltage drop over a calibrated shunt resistor, using a digital oscilloscope (Picoscope 2206D, input impedance 1 MΩ). All currents reported in this paper are these measured current values. The stimulation experiments were controlled by interfacing the function generator with a Python code to run the experimental sequence allowing the trial parameters to be randomized within an experimental block and ramping procedure to be consistent across trials.

### Stimulation experiments on *Locusta migratoria*

Animals were obtained at a local pet shop and kept in the laboratory in a terrarium with heat lamp with grass and oats *ad libitum*. The only inclusion criteria for performing the experiment were adult individuals with fully intact anatomy and lack of any obvious superficial injuries. Prior to the stimulation experiment, we anesthetized the locust by cooling to 5 °C for around 15 min. Next, the animal was immobilized with the ventral side facing upward, in a bed made of modelling clay, and allowed to warm up to room temperature prior to starting the experiment. Stimulation was delivered via 2 or 4 Ag/AgCl wires placed onto the cuticle using *xyz* micromanipulators (S-725CRM, Signatone USA). Contact between the wire and the cuticle was enhanced by adding a drop of EKG gel onto the tip of the wire prior to making contact with the cuticle. In the optimization of the 4-electrode TIS experiment, we used an oscilloscope and microwire probe, relative to a ground electrode in the tail of the animal, to verify the “hot spot” of high amplitude modulation depth and high relative amplitude. The video of the leg movement was recorded using a digital camera (Sony DSC-RX II) with a frame rate of 100 fps. An initial threshold at AMF=1 Hz was obtained to detect the minimum detectable leg twitch, approx. 5 degrees. Stimulation was started and AMF frequency changed once 5 kicks were observed at a given frequency. Deflection angles as a function of AMF were obtained using a free and open-source video analysis tool *Tracker* (https://opensourcephysics.github.io/tracker-website/). Temporal syncing of the video to the stimulation waveform was done by triggering an LED which produced a light impulse (visible in one frame of video) at the beginning of the waveform and again at the AM maximum of the waveform. Data were obtained from naïve animals, and individuals were not reused for subsequent experiments with data collection. We would note, however, that the animals survive this experimental sequence without a problem and can be returned to the terrarium after marking with a colored permanent marker. The locust experiment to determine the s-f curve was coded in such a way that the presentation of a given frequency was randomized within a waveform block. A 30 second to 1-minute gap was taken in between each trial. The exact location of the two-electrode stimulation was always determined from the position of the TIS electrodes, therefore the block of TIS stimulation always came first. After this the choice of waveforms was randomized. Ramping to threshold was carried out identically for each waveform. In all locust experiments the ramping procedure was as follows: The Keysight was set to 0.005 V_pp_ and stimulation started. Every 0.5 sec the amplitude was increased by 0.005 Vpp until the threshold was observed. At this point the amplitude was decreased by 0.005 Vpp every 0.5 sec until the evoked activity disappeared, then the amplitude was again increased by 0.005 Vpp until the evoked action reappeared. To ensure an approximately equal time for each trial, frequencies >5 kHz had a max output current set to 10 mA, trials with frequencies < 5 kHz had a max output current set to 1 mA.. Each trial began again from the same initial amplitude value. For experiments with frequencies >12.5kHz an automated ramping procedure could not be used due to the increased variability in the ability to stimulate the N5 nerve. As such manual ramping was used so that the threshold values could precisely be determined. Similarly, because there is a higher likelihood of damaging the ability of the leg to kick at higher frequencies, randomization of frequency presentation was not possible. Instead, frequencies were applied in increasing order.

### Stimulation experiments on human subjects

Subjects were healthy volunteers who expressed written informed consent. To ensure pressure exerted on the forearm, and thus movement on the muscles during the experiment, was minimized the participants hand was placed on a cushion with their elbow on the table. This meant the forearm was not in contact with the table and a more stable position obtained. Electrodes were placed onto the skin and adhered in place using tape. The primary stimulation electrodes were always gel-assisted AgCl EKG electrodes, (Ceracarta), in two-electrode stimulation protocols the ground electrode was a larger TENS pad electrode 4 × 4 cm^2^. Subjects were not told of the specific sequence of stimulation or variable changes but were not otherwise blinded. The experiment was coded in such a way that the presentation of a given frequency was randomized within a waveform block. A 30 second to 1-minute gap was taken in between each trial. The exact location of the two-electrode stimulation was always determined from the position of the TIS electrodes, therefore the block of TIS stimulation always came first. After this the choice of waveforms was randomized. Ramping to threshold was carried out identically for each waveform. In all human experiments the ramping procedure was as follows: The Keysight was set to 0.05 Vpp and stimulation started. Every 0.5 sec the amplitude was increased by 0.05 Vpp until the threshold was observed (or reported by the participant). At this point the amplitude was decreased by 0.05 Vpp every 0.5 sec until the evoked activity disappeared, then the amplitude was again increased by 0.05 Vpp until the evoked action reappeared. To ensure an approximately equal time for each trial frequencies < 5kHz had an incrementation rate of 0.005 Vpp. Each trial began again from the same initial amplitude value. This general scheme was applied to all human experiments.

#### Participants

Participants were recruited via an advertisement in the local community and on the institute’s campus. A total of N=8 participants (average age 34.25 ± 11.2, 62% Female) were recruited for the motor stimulation of the median nerve. For our afferent nerve stimulation experiments N=11 participants (average age = 36.3 ± 8.21, 36% Female) were recruited. For the determination of the ‘U-curve’ in the afferent model N=6 participants were recruited (average age = 33 ± 10, 83% Female); for the efferent model N=7 participants (average age = 27 ± 5, 28% female)

### Computational modeling

NEURON model (used for Figure 1d) was taken from Mirzakhalili et al.,^17^ the model files are published and freely available online at https://github.com/mirzakhalili/Mirzakhalili-et-al<u>–CellSystems-2020</u>. The computation for Figure 1d was made by simulating 1 kHz extracellular axonal stimulation for two different current amplitudes 2 mA (subthreshold) and 3 mA (suprathreshold). Sinusoidal stimulation was obtained by setting Δ𝑓 = 0 Hz and TIS stimulation was obtained by setting Δ𝑓 = 5 Hz. The results were evaluated at node 45.

Finite-element modeling (Figures 4 and 5): Calculations were done in COMSOL Multiphysics 6.3 using the AC/DC Module and Electric currents physics interface. The study was solved in frequency domain for 2500 Hz carrier frequency. The forearm is implemented as a three-layer model inspired by the work by Medina and Grill where the layers represent skin, fat and muscle.^7^ Between the skin and the gelled ECG electrodes is a distributed electrode-skin interface (ESI) impedance consisting of a constant phase element CPE with parameters 𝐾 = 35 MΩs^−α^cm^2^ and 𝛼 = 0.9 in parallel to a large charge transfer resistance 𝑅𝑝 = 4.7 MΩcm^2^ with the parameter values as reported by McAdams et al.^64^ The boundary condition on the ESI is given by 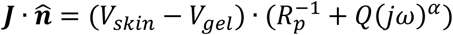, where 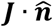 is normal current density at the interface elements, *V_skin_* and *V_gel_* are the electric potentials on each side of the ESI, while the second bracket is the admittance of the CPE. The boundary condition on the electrodes is set by the total current *I_tot_* flowing through the electrode surface 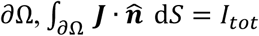. We perform two independent simulations to calculate the current contributions from each electrode pair, in one simulation we set *I_tot_* = ±𝐼_1_ on the first pair of electrodes and *I_tot_* = 0 for the second pair, and in the second simulation *I_tot_* = 0 for the first pair and *I_tot_* = ±𝐼_2_ for the second pair, finally, the total current is obtained by adding the contributions from each pair. It is important to do the simulation in this way because the current stimulators used in the experiment are isolated. All other external boundaries have electrical insulation boundary condition 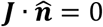. In the bulk the current continuity equation -∇(𝜎𝛁𝑉 + 𝜖_0_𝜖𝑟𝑗*ω*𝛁𝑉) = 0 is solved for electric potential 𝑉, where the first contribution is from the Ohmic current parameterized by bulk conductivity 𝜎 and the second contribution is displacement current parameterized by relative permittivity 𝜖𝑟. For the muscle tissues 𝜎 and 𝜖𝑟 are anisotropic with different values along the muscle fibers (x-direction) and across the muscle fibers (y,z direction). The entire geometry is meshed with a tetrahedral mesh with 1.2 · 10^6^ domain elements with sizes ranging from 0.5 mm near the surface layers to 5 mm deep in the bulk. Activating functions are calculated as second-order spatial derivatives of the electric potential in the nerve direction (*x*-direction). Values of the material parameter used in the simulation for the frequency 2.5 kHz are listed in Table 1 are taken and adapted from Gabriel et al^65^.

| Material | Conductivity $\sigma$ [S/m] | Relative permittivity $\epsilon_r$ |
| --- | --- | --- |
| Muscle along | 0.6 | $1.5 \cdot 10^5$ |
| Muscle across | 0.35 | $1 \cdot 10^5$ |
| Adipose tissue (fat) | 0.024 | 3000 |
| Skin | 0.0015 | 40000 |
| ECG gel | 0.2 | 70 |

## Author contributions

A.O. conducted all computational modeling, with support from V. Đ. P.O. and E.D.G., supported by J.T., performed all preliminary experiments on TIS comparisons with kHz in locusts and humans, and P.O. made the first observation that thresholds of kHz stimulation appear similar for modulated and unmodulated waveforms. A.O, and D.S.R. performed all final locust experiments and data analysis. Final human data was collected and analyzed by D.S.R. V.Đ. and E.D.G. supervised the work and obtained funding. A.O., V.Đ., and E.D.G. conceived the project idea and performed conceptualization. The manuscript was written by E.D.G. with input from all authors.

